# Disease-associated mutations in the STAT5B SH2 domain reprogram hepatic cholesterol and lipid metabolism

**DOI:** 10.1101/2025.11.27.691004

**Authors:** Hye Kyung Lee, Maxim Pyatkov, Oksana Gavrilova, Naili Liu, Tamar Demby, Bingtian Ye, Priscilla A. Furth, Lothar Hennighausen, David J Waxman

**Author notes:** Correspondence to: L.H, D.J.W.

## Abstract

Growth hormone (GH) signaling through STAT5B is a central regulator of hepatic metabolism, yet the functional consequences of disease-associated STAT5B variants remain poorly understood. Here, we analyzed mice carrying STAT5B^Y665F^ (gain-of-function) and STAT5B^Y665H^ (loss-of-function) variants and dissect their impact on metabolic regulation. STAT5B^Y665F^ mice developed hepatic lipid accumulation, hypercholesterolemia, and enhanced insulin sensitivity, whereas STAT5B^Y665H^ mice displayed reduced body weight and impaired insulin responsiveness. Transcriptomic analyses revealed that STAT5B^Y665F^ activated lipid, cholesterol, and immune transcriptional programs, while STAT5B^Y665H^ failed to induce these pathways. Notably, STAT5B^Y665F^ substantially feminized male liver gene expression, inducing 77% of female-biased genes while repressing 51% of male-biased genes, thereby mimicking the persistent STAT5B activation characteristic of female livers. ChIP-seq demonstrated extensive STAT5B^Y665F^ enhancer occupancy at metabolic and immune loci, contrasting with the minimal chromatin engagement of STAT5B^Y665H^. Beyond the liver, STAT5B^Y665F^ broadly reprogrammed adipose tissue gene expression, activating lipid metabolism and immune regulatory networks, whereas STAT5B^Y665H^ exerted more restricted effects. Together, these findings illustrate how alterations in STAT5B activity affect enhancer activation and can lead to changes in metabolic function and hepatic sexual dimorphism.

## Introduction

The Signal Transducer and Activator of Transcription 5 (STAT5) is a key transcription factor that integrates hormonal signals to regulate growth, immune function, and metabolism (1). Among its isoforms, STAT5B is predominant in the liver, where it mediates growth hormone (GH) signaling (2–4) and controls metabolic pathways involved in lipid metabolism, glucose homeostasis, and cholesterol biosynthesis (5–9). STAT5B serves as a critical transcriptional regulator of genes that maintain cholesterol balance and lipid utilization, thereby contributing to overall metabolic stability.

STAT5B is essential for GH-mediated activation of genes such as insulin-like growth factor 1 (IGF-1) (2,3,10,11) and for modulating networks involving peroxisome proliferator-activated receptors (PPARs) (12,13). It also influences sex-dependent liver gene expression (14–16), impacting hepatic metabolism and highlighting how its dysregulation can contribute to metabolic disorders. Dysregulated STAT5B signaling has been implicated in insulin resistance, metabolic dysfunction-associated steatotic liver disease (MASLD), obesity and dyslipidemia (17–19), and is associated with worsened liver injury and metabolic decline (20,21). For example, Río-Moreno et al. (20) showed that impaired hepatocyte GH/STAT5B signaling exacerbates liver injury in MASLD models, while Davey et al. (21) demonstrated that STAT5B-deficient mice are resistant to GH pulses characteristic of male liver (22), underscoring its role in maintaining normal metabolic responses. Beyond metabolism, STAT5B influences inflammatory and fibrogenic processes (8,23), as well as the hepatic chromatin landscape (24,25), further linking it to liver disease progression.

The most frequent STAT5B variants, N642H and Y665F, both enhance STAT5 activity (26,27). Within the SH2 domain, tyrosine-665 (Y665) can be substituted with phenylalanine (Y665F) or histidine (Y665H) (28,29). While Y665F is recurrently observed in patients with hematologic diseases, such as Large Granular Leukemia, (28–30), Y665H has been reported only once, in a T-prolymphocytic leukemia case (28). STAT5^Y665^ variants show Gain-of-Function (GOF) activity (31) *in vitro*, suggesting they may have overlapping transcriptional effects. However, *in vivo* studies reveal that STAT5B^Y665F^ retains GOF activity, whereas STAT5B^Y665H^ has characteristics of a Loss-of-Function (LOF) variant (32–34), and neither mutation alone induces malignant transformation. Thus, these STAT5^Y665^ substitutions can differentially alter STAT5B signaling to downstream pathways.

Tyrosine 699 serves as the critical JAK2 phosphorylation site (29,33) and tyrosine 665 contributes to STAT5B dimerization, nuclear translocation, and downstream transcriptional regulation (32,33). Our previous studies have shown that these mutations produce tissue-specific effects: in the mammary gland, Y665F causes hyperproliferation and increased the expression of milk protein genes (GOF phenotype), whereas Y665H results in developmental delay and reduced proliferation (LOF phenotype) (33). In immune cells, Y665F drives T-cell hyperproliferation and inflammatory responses, while Y665H impairs T-cell differentiation (32).

The role of STAT5B in sexual dimorphism adds another layer of complexity, as GH signaling exhibits sex-dependent dynamics that influence hepatic metabolism (16,22,35–37) and liver disease susceptibility (38). STAT5B mutations may therefore differentially affect males and females, producing sex-specific alterations in metabolic homeostasis and disease.

Given these profound, context-dependent effects, we hypothesize that STAT5B^Y665F^ and STAT5B^Y665H^ variants will exert distinct influences on hepatic metabolism. In this study, we combine genomic and functional analyses to investigate their impact on cholesterol homeostasis and lipid metabolism *in vivo*. These findings help clarify the mechanistic basis of STAT5B-mediated metabolic regulation and may inform targeted therapeutic strategies for STAT5B-related metabolic disorders, including MASLD and obesity.

## Results

### Liver metabolic phenotype in STAT5B^Y665F^ and STAT5B^Y665H^ mutant mice

To elucidate how STAT5B^Y665F^ and STAT5B^Y665H^ mutations affect GH-regulated metabolic pathways, we examined mice carrying these variants (32,33). To investigate potential molecular defects arising from these mutations, we performed histological and metabolic analyses on two-month-old male and female mice. H&E-staining revealed that, in both sexes, STAT5B^Y665F^ livers contained more unstained (white) areas within hepatocytes, indicative of lipid accumulation, whereas STAT5B^Y665H^ livers resembled wild type (WT) (Figure 1A, Supplementary Figure 1A).

**Figure 1.**
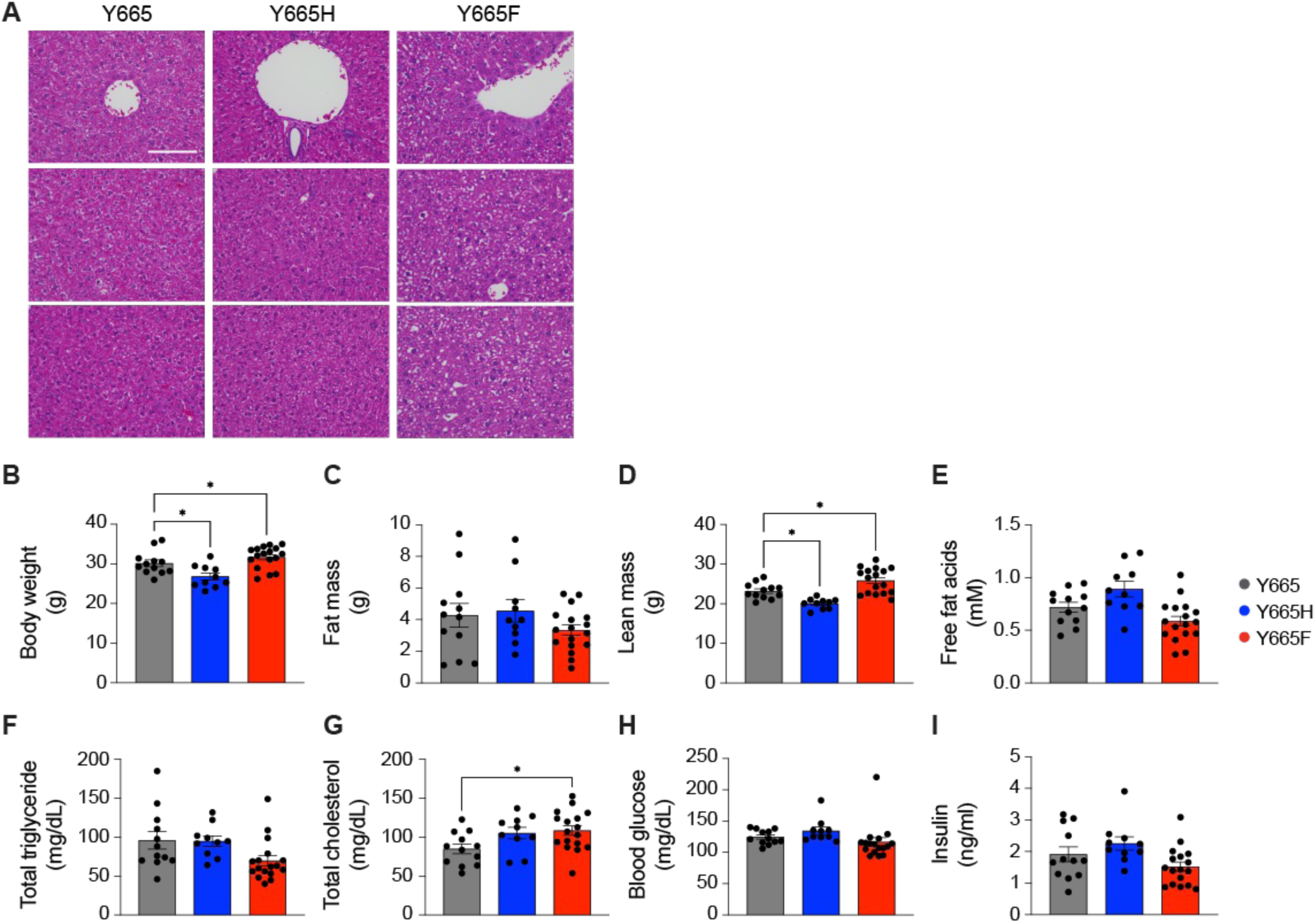
Metabolic phenotypes regulated by STAT5B^Y665^ mutations. **(A)** Representative hematoxylin and eosin (H&E)-stained liver sections from 2-month-old male mice (magnification x400; scale bars, 100 μm). **(B-I)** Body weight (B), plasma fat mass and lean mass (C-D), free fat acids (E), total triglyceride (F), total blood cholesterol (G), blood glucose (H), and insulin (I) levels in WT and mutant male mice at 14-15 weeks of age. Data are presented as the mean ± SEM for independent biological replicates (WT, *n* = 12; Y665H, *n* = 10; Y665F, *n* = 17). *P*-value were determined by one-way ANOVA with Tukey’s multiple comparisons test. \**p* < 0.05.

We next assessed systemic metabolic parameters, including body weight, fat/lean mass, free fatty acids, triglycerides, cholesterol, blood glucose and insulin levels in male mice (Figure 1B-I, Supplementary Figure 1B-I). Because many GH-regulated phenotypes are mediated by insulin-like growth factor 1 (IGF-1), we measured serum IGF-1 and hepatic *Igf1* mRNA levels. Only STAT5B^Y665F^ male mice showed significant increases compared to WT mice, whereas STAT5B^Y665H^ mice did not differ from WT mice (Supplementary Figure 2A-B). STAT5B^Y665H^ male mice exhibited significantly reduced body weights compared to WT, while STAT5B^Y665F^ mutants were significantly heavier (Figure 1B). At 14 weeks of age, STAT5B^Y665F^ mice tended to have lower fat mass, free fatty acids, and triglycerides, though the differences were not statistically significant (Figure 1C, E, F). Lean mass was significantly increased in STAT5B^Y665F^ but reduced in STAT5B^Y665H^ mice (Figure 1D). Importantly, plasma cholesterol levels were significantly elevated in STAT5B^Y665F^ mice (Figure 1G), indicating a gain-of-function (GOF) effect on cholesterol metabolism that is dependent of changes in IGF-1. Blood glucose and insulin levels showed modest, non-significant increases in STAT5B^Y665H^ and decreases in STAT5B^Y665F^ males (Figure 1H-I). Similar to males, STAT5B^Y665F^ females displayed elevated cholesterol (Supplementary Figure 1G), whereas STAT5B^Y665H^ females had significantly higher insulin levels (Supplementary Figure 1I). Together, these findings demonstrate that STAT5B^Y665F^ and STAT5B^Y665H^ variants exert opposing effects on liver metabolism: Y665F drives lipid accumulation and hypercholesterolemia, while Y665H confers a leaner phenotype with relatively preserved metabolic function.

### Glucose metabolism in STAT5B^Y665^ variant mice

To assess glucose homeostasis, we performed a homeostatic model assessment for insulin resistance (HOMA-IR) and assayed fasting blood glucose and fasting insulin levels in 14-15-week-old male mice. Although no statistically significant differences were detected, values tended to be higher in STAT5B^Y665H^ males and lower in STAT5B^Y665F^ males (Figure 2A-C).

**Figure 2.**
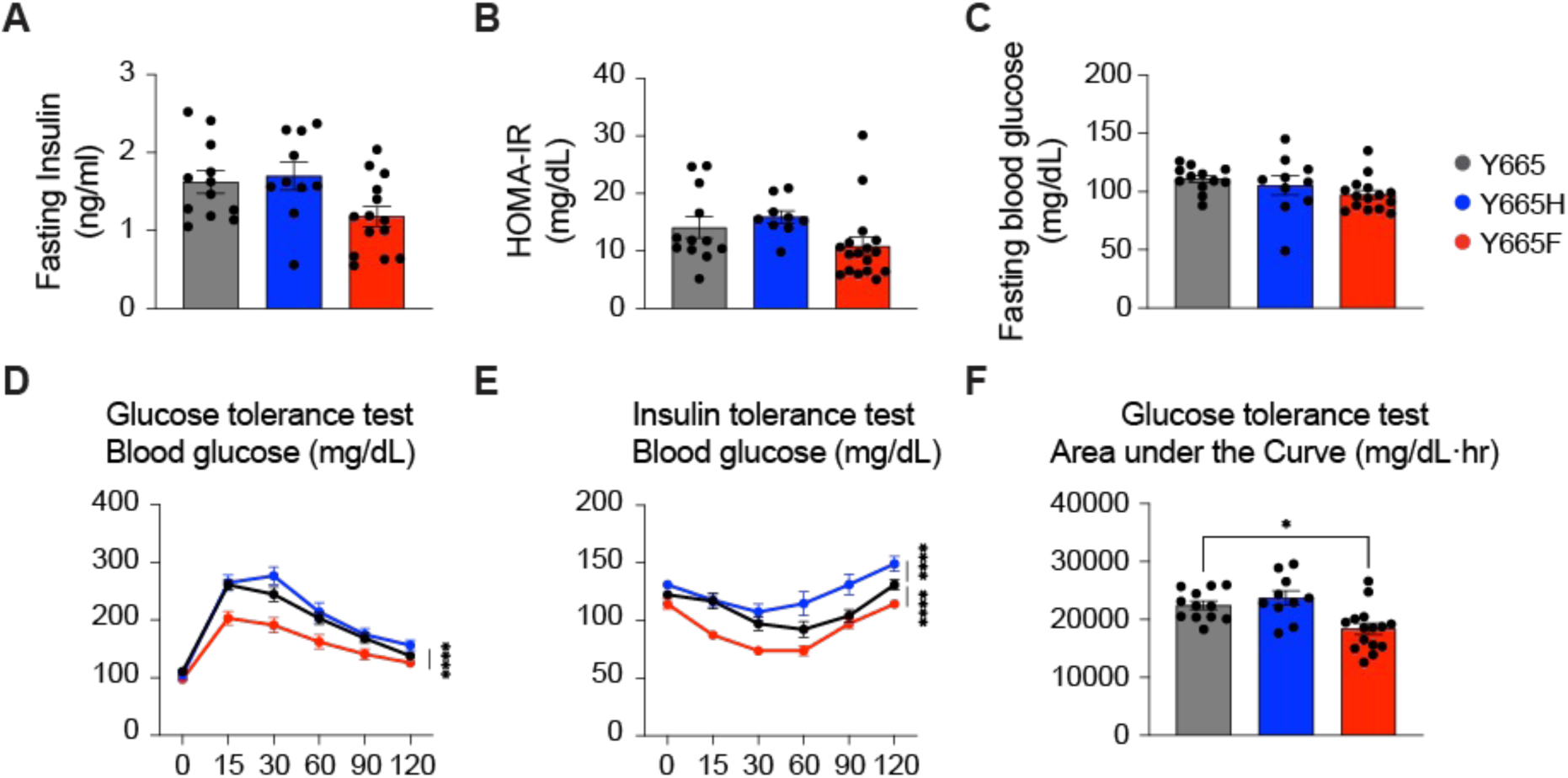
Glucose tolerance and insulin sensitivity in STAT5B^Y665^ mutant mice. **(A-B)** Fasting blood glucose (A) and fasting insulin (B) levels. Data are shown as the mean ± SEM (WT, *n* = 12; Y665H, *n* = 10; Y665F, *n* = 17). *P*-value were calculated using one-way ANOVA with Tukey’s multiple comparisons test. **(C)** Homeostatic Model Assessment for Insulin Resistance (HOMA-IR) levels. **(D-E)** Glucose tolerance test (GTT) and insulin tolerance test (ITT) in 14-15-week-old WT and STAT5B^Y665^ male mutant mice. Line graphs show blood glucose concentrations up to 120 min after glucose or insulin administration. *P*-value were determined by two-way ANOVA with Tukey’s multiple comparisons test. \*\*\*\**p* < 0.0001. **(F)** Area under the curve (ACU) of GTT, reflecting total blood glucose following glucose load. \**p* < 0.05.

We then performed glucose tolerance tests (GTT) and insulin tolerance tests (ITT). ITT was performed five days after GTT following a 2-hour fast (Figure 2D-E). In both tests, STAT5B^Y665F^ mice displayed significantly lower blood glucose levels and a stronger reduction after insulin injection, consistent with enhanced insulin sensitivity (Figure 2D-E). In contrast, STAT5B^Y665H^ mice showed mildly elevated fasting glucose and insulin values, but these differences were not statistically significant; their glucose clearance during GTT resembled WT, while ITT responses were impaired (Figure 2D-E). The area under the curve (AUC) for GTT was significantly reduced in STAT5B^Y665F^ mice, reflecting improved glucose clearance (Figure 2F). Similar trends were observed in female mice (Supplementary Figure 1L-N). These findings indicate that STAT5B^Y665F^ enhances insulin sensitivity and glucose clearance, suggesting a protective metabolic effect, whereas STAT5B^Y665H^ does not confer such benefits and instead shows signs of impaired insulin responsiveness.

### Transcriptional impacts of STAT5B^Y665^ variants in liver

To investigate transcriptional consequences of each STAT5B^Y665^ variant, we conducted transcriptomic (RNA-seq) and chromatin immunoprecipitation sequencing (ChIP-seq) analyses on mouse liver tissue. In male liver, STAT5B^Y665F^ uniquely altered the expression of 1,261 genes, while STAT5B^Y665H^ uniquely dysregulated 115 genes, and 134 genes showed a common response to both STAT5^Y665^ mutations when compared to WT livers (Figure 3A; Supplementary Table 1). Pathway analysis of significantly regulated genes from both groups revealed enrichment in retinol, lipid, and cholesterol metabolism, as well as inflammatory response pathways (Figure 3B). Both the up- and down-regulated genes from the two variants were analyzed (Figure 3C). Notably, cytokine production and leukocyte activation pathways were selectively enriched in STAT5B^Y665F^ livers. Genes involved in cholesterol/lipid metabolism and cytokine signaling were strongly upregulated in livers of STAT5B^Y665F^ mice but reduced or only weakly induced in STAT5B^Y665H^ mice (Figure 3C), consistent with the elevated blood cholesterol levels seen in STAT5B^Y665F^ livers (Figure 1G) and with the activation of cholesterol-regulatory gene networks including cholesterol metabolism enzymes (Supplementary Figure 3C-D). Because hepatic cholesterol synthesis and uptake pathways are primary determinants of whole-body cholesterol homeostasis, altered expression of cholesterol-metabolizing enzymes can directly influence circulating cholesterol levels (39,40). Thus, the coordinated upregulation of these enzymes in STAT5B^Y665F^ livers provides a mechanistic explanation for the increased serum cholesterol observed in these mice.

**Figure 3.**
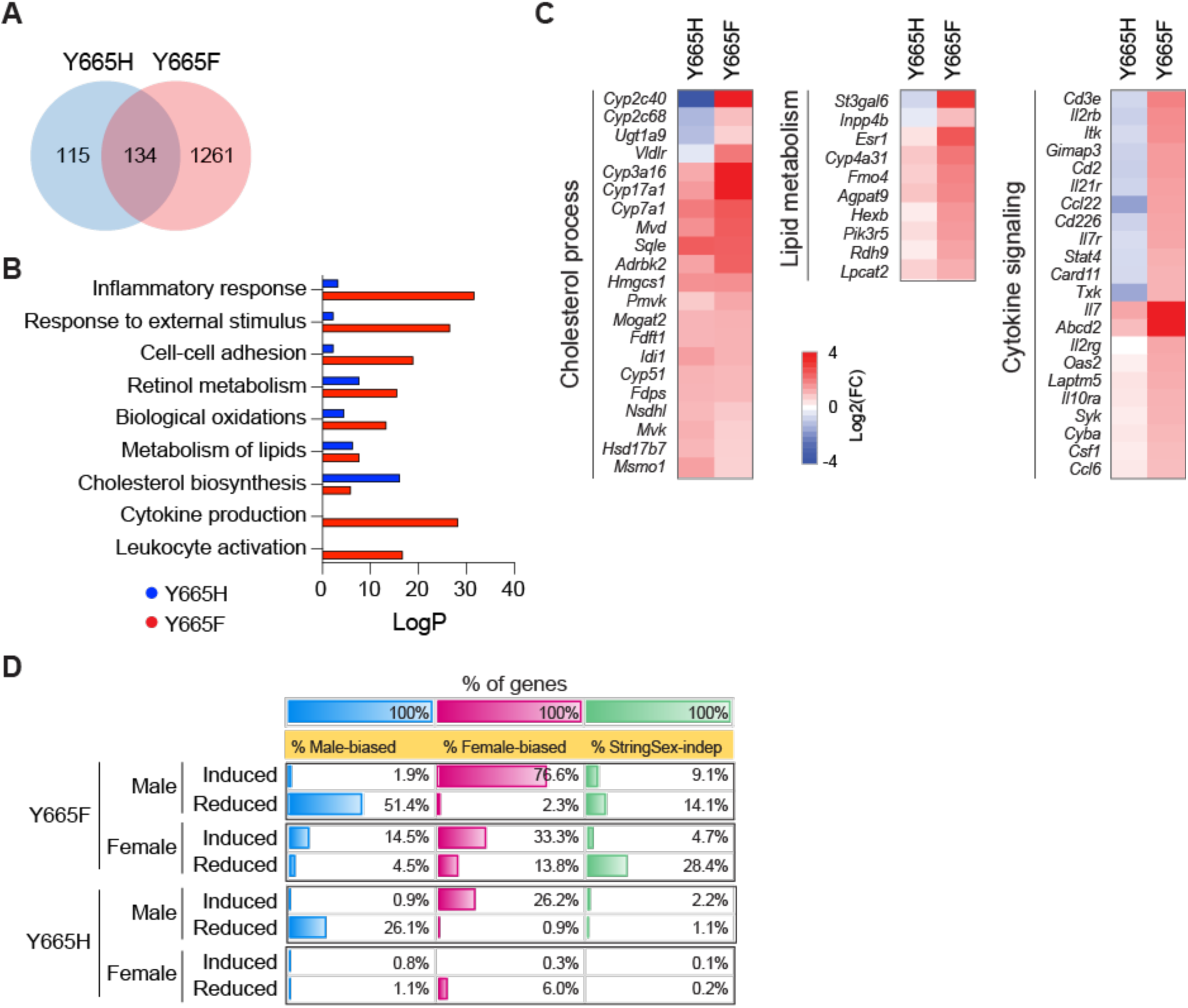
Divergent transcriptional activity of STAT5B^Y665F^ and STAT5B^Y665H^ in liver. **(A)** Venn diagram displaying the number of significantly regulated genes in STAT5B^Y665^ mutant male mice compared to WT, as evaluated by RNA-seq (*n* = 6). **(B)** Bar graph of significantly differentially expressed genes (DEGs) enriched in Gene Ontology (GO) terms. **(C)** Heatmaps of log2(fold changes) in DEGs related to cholesterol/lipid metabolism and cytokine signaling. Red, indicates genes induced compared to WT and blue indicates genes whose expression is reduced compared to WT. **(D)** Patterns of response to STAT5B^Y665F^ and STAT5B^Y665H^ in male liver and in female liver, as marked, for the sets of genes showing significant differences in expression between WT male and female liver, and for a set of stringently sex-independent genes (see Methods).

In females, transcriptional changes were less extensive but revealed distinct regulatory features. Both the up- and down-regulated genes from the two variants were analyzed. RNA-seq identified 46 genes commonly dysregulated in both mutants compared to WT, with 781 genes uniquely regulated in STAT5B^Y665F^ and 88 genes uniquely responsive to STAT5B^Y665H^ (Supplementary Figure 3A; Supplementary Table 2). Pathway analysis of significantly regulated genes from both groups indicated increases in lipid and cholesterol metabolism and cytokine signaling (Supplementary Figure 3B). Unlike in males, where broad immune pathways were strongly activated, female STAT5B^Y665F^ mutants displayed a narrower but focused set of immune-related differentially expressed genes (DEGs) (Supplementary Figure 3C), suggesting sex-specific modulation of hepatic immunity. These transcriptional changes are consistent with the robust increase in blood cholesterol seen in STAT5B^Y665F^ females (Supplementary Figure 1G), though the overall magnitude of metabolic reprogramming was lower than in males. Thus, STAT5B^Y665F^ broadly activates lipid, cholesterol, and immune transcriptional programs, whereas STAT5B^Y665H^ fails to induce these pathways. In females, the transcriptional effects of STAT5B^Y665F^ are more limited, revealing sex-dependent differences in STAT5B-mediated metabolic gene regulation.

Comparison of the transcriptional changes induced by STAT5B^Y665F^ to the genes showing significant male-female differences in expression in wild-type (WT) mouse liver revealed the following (Figure 3D, Supplementary Table 3). In male liver, STAT5B^Y665F^ induced 77% of the genes that showed higher expression in WT female than WT male liver (i.e., female-biased genes), whereas it repressed only 2.3% of these genes. In contrast, STAT5B^Y665F^ repressed 51% of the genes showing higher expression in WT male than WT female liver (male-biased genes), whereas it induced only 1.9% of these genes. Thus, STAT5B^Y665F^ substantially feminizes liver gene expression in male liver. STAT5B^Y665H^ similar trends, albeit affecting only 26% of male-biased and female-biased genes in male liver, consistent with the more limited actions of STAT5B^Y665H^ described above. Hierarchical clustering of WT liver sex-biased genes validated these patterns and confirmed the co-clustering of gene expression patterns for male-STAT5B^Y665F^ with female-WT livers, as well as co-clustering for male-STAT5B^Y665H^ livers with male-WT livers (Figure 4A, *top*). These STAT5-Y655 variant response patterns were not apparent in female liver (Figure 3D, Figure 4B) and were also not seen when we examined gene responses in a set of 3,414 stringently sex-independent genes (Figure 3D). In male liver, STAT5B^Y665F^ induced many female-biased genes to 100% of WT female liver levels, with supraphysiological levels of expression (>2-fold above WT female levels) observed for 27% of female-biased genes (Supplementary Table 3A; e.g., Figure 4A, cluster 5). STAT5B^Y665F^ also induced supraphysiological expression of many female-biased genes in female liver (e.g., Figure 4B, cluster 3). Six highly female-biased genes were induced by STAT5B^Y665F^ to levels ∼10 to 25-fold higher than in WT female liver, as seen in both male and female liver: *A1bg, Cyp3a41a, Cyp3a41b, Sult2a1, Sult2a3* and *Sult3a2* (Supplementary Table 3A).

**Figure 4.**
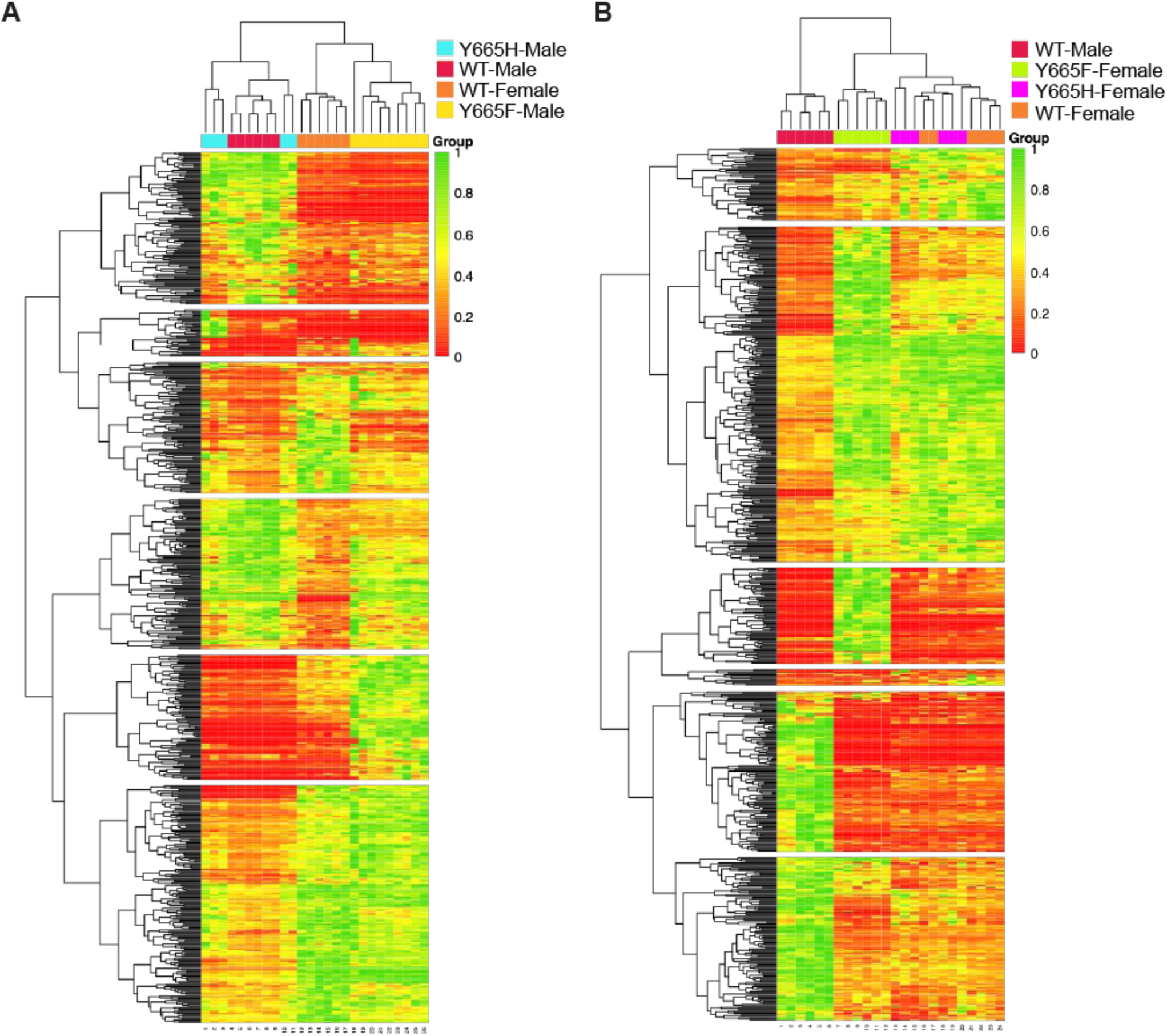
Heatmap showing response patterns to STAT5B-Y665 mutations in male (A) and in female liver (B) for those genes showing significant expression differences between male and female mouse liver. Hierarchical clustering was applied to sex-biased genes meeting these thresholds: WT male/female |fold-change| > 2, padj < 0.05, and TPM > 1, and excluding all genes with lncRNA annotations (n = 664 genes). Each column represents RNA-seq profile for one individual mouse from each of the 4 groups shown. Normalized gene expression is scaled from 0 (*red*, lowest expression across all livers) to 1 (*green*, highest expression). Gene clusters are numbered from top to bottom and are detailed in Supplemental tables 3C and 3D.

Comparison with other STAT5 models further underscores the opposing activities of the Y665 variants. STAT5B^Y665F^ shared 361 upregulated genes with constitutively active STAT5 (STAT5B_CA_) (41), which were enriched in immune and inflammatory pathways such as leukocyte activation, cytokine production, and antiviral responses, as well as lipid-related processes including retinol, arachidonic acid, and peroxisomal metabolism (Supplementary Table 1, Supplementary Table 2). This overlap indicates that STAT5B^Y665F^ recapitulates key features of constitutive STAT5 activation and is consistent with the pattern of feminization by STAT5B^Y665F^ described above. Conversely, STAT5B^Y665H^ showed transcriptomic convergence with STAT5-deficient livers (STAT5KO) (14), with 44 genes significantly downregulated in both models in males—linked to retinol metabolism, lipid oxidation, and organic acid catabolism—and 42 genes suppressed in females, associated with steroid biosynthesis, glucose metabolism, and adipocyte differentiation (Supplementary Table 1, Supplementary Table 2). These comparisons indicate that STAT5B^Y665F^ resembles a constitutively hyperactive STAT5 state, which may mimic the near persistent activation by GH of STAT5B signaling seen in female liver (41), whereas STAT5B^Y665H^ approximates partial loss of function, highlighting Y665 as a molecular switch modulating STAT5B activity across metabolic and immune networks.

### Chromatin regulation by STAT5B^Y665F^ and STAT5B^Y665H^

ChIP-seq analysis revealed marked differences in STAT5B chromatin binding. In STAT5B^Y665F^ livers, strong STAT5B binding peaks co-localized with H3K27ac and RNA polymerase II (Pol II), consistent with active enhancer engagement. In contrast, STAT5B^Y665H^ showed markedly reduced STAT5B occupancy and diminished liver chromatin accessibility (Figure 5; Supplementary Table 4). Motif and chromatin mark profiling further underscored these differences (Figure 5A). In WT livers, 10,753 STAT5B peaks were detected, of which approximately 30% (3,190/10,753) contained a canonical GAS motif (TTCnnnGAA) (Table S4A), of which 5% (161/3,190) overlapped with H3K27ac (Table S4B). STAT5B^Y665H^ mutants exhibited only 143 peaks, with only 4 (2.8%) containing a GAS motif, of which 2 overlapped with H3K27ac (Table S4C, Table S4D). By contrast, STAT5B^Y665F^ mutants displayed 13,477 peaks, with 24% (3,277/13,477) harboring a GAS motif (Table S4E), of which 21% (701/3,277) co-localized with H3K27ac (Table S4F). Among these, 701 peaks contained both a GAS motif and H3K27ac marks, with 54% (intergenic 169 + intronic 209 / 701) located in intergenic or intronic regions, consistent with putative enhancer elements (Figure 5B). These enhancer-associated peaks were enriched near genes involved in immune regulation, adipogenesis, and hepatic metabolic processes (Figure 5C). Of the 1,395 genes transcriptionally activated in STAT5B^Y665F^ livers, a subset was directly bound by STAT5B (Supplementary Table 4). Canonical STAT5 targets, such as *Cish,* were induced, and key cholesterol metabolism genes (*Cyp51*, *Esr1*, *Fdft1*, *Lpin1*, *Fut8*) were upregulated in STAT5B^Y665F^ livers, whereas these pathways remained inactive in STAT5B^Y665H^ livers (Figure 5D). In females, STAT5B^Y665F^ mutants exhibited 25,965 peaks, with 21% (5,459/25,965) harboring a GAS motif, of which 34% (1,836/5,459) co-localized with H3K27ac. Among these, 1,836 peaks carried both a GAS motif and H3K27ac marks, with 59% (intergenic 441 + intronic 650 /1,836) located in intergenic or intronic regions (Supplementary Figure 4A-C; Supplementary Table 5). Together, these results show that STAT5B^Y665F^ establishes an active enhancer landscape at many loci, including key metabolic and immune loci, whereas STAT5B^Y665H^ fails to maintain WT STAT5B levels of chromatin accessibility, highlighting the fundamentally opposing roles of these two mutants in transcriptional regulation.

**Figure 5.**
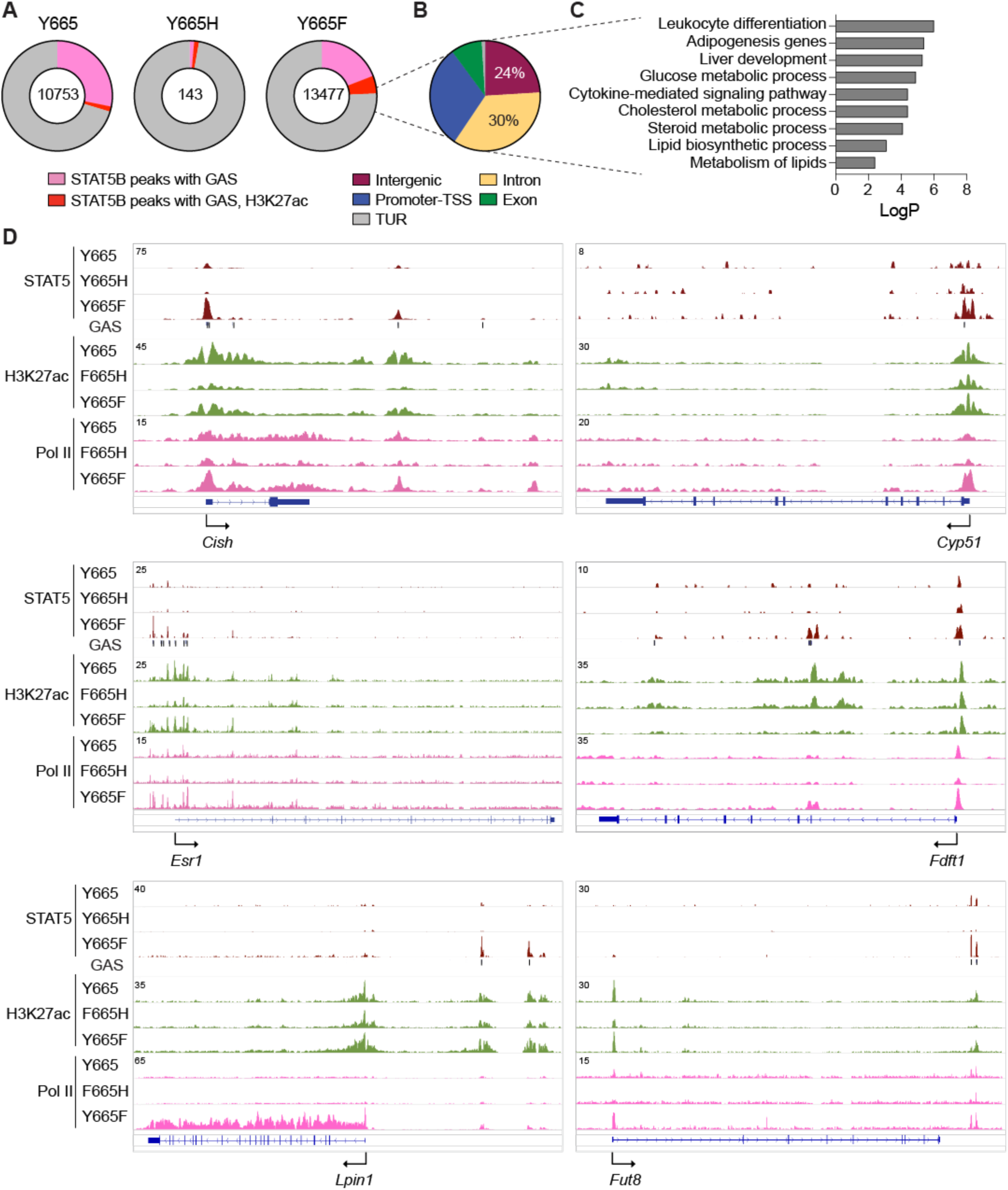
STAT5B^Y665F^ and STAT5B^Y665H^ exert opposing effects on enhancer activation in liver. **(A-B)** Distribution of STAT5B binding peaks containing GAS motifs and overlapping with H3K27ac marks. Pie charts show ChIP-seq signals within ±3 kb of STAT5B binding sites in WT and mutant male livers. **(C)** Bar graph of target genes associated with enhancer clusters enriched in GO terms. **(D)** Genome browser tracks showing STAT5B binding, H3K27ac and Pol II occupancy at regulatory elements of Cytokine-inducible SH2-containing protein (*Cish*), Cytochrome P450 family 51 (*Cyp51*), Estrogen receptor 1 (*Esr1*), Farnesyl-Diphosphate Farnesyltransferase 1 (*Fdft1*), Lipin1 (*Lpin1*), Fucosyltransferase 8 (*Fut8*) and in liver tissue from WT, STAT5B^Y665H^ and STAT5B^Y665F^ mice.

### Contrasting transcriptional programs driven by STAT5B^Y665F^ and STAT5B^Y665H^ in lipid metabolism

Since hepatic lipid metabolism influences adipocyte function (42), we investigated the impact of STAT5B mutations on lipid-associated transcriptional programs in adipose tissue. Both the up- and down-regulated genes from the two variants were analyzed. In male mice, RNA-seq profiling identified 670 genes commonly dysregulated by both STAT5B mutations relative to WT, with 424 genes uniquely altered in STAT5B^Y665H^ and 680 uniquely altered in STAT5B^Y665F^ mice (Figure 6A; Supplementary Table 6). Gene ontology (GO) analysis of significantly regulated genes from both groups revealed significant enrichment for pathways related to male meiotic nuclear division, ion transport, glycerophospholipid biosynthesis, and cytokine-mediated immune responses (Figure 6B). DEGs included cytokine/immune regulators (*Gzmb*, *Ccr3*, *Il2rb*, *Prf1*, *Lat*, *Camk4, Sla2, Itga2, Relt*), lipid metabolism genes (*B3galt5*, *F7, Plcb2*, *Cav3*), and ion transport components (*Kcnc1*, *Gp9*, *Gimap3*, *Cracr2a, Ccl5, Cxcr3*) (Figure 6C). In females, RNA-seq profiling identified 68 commonly dysregulated genes, with 245 uniquely altered in STAT5B^Y665H^ and 667 uniquely altered in STAT5B^Y665F^ mice (Supplementary Figure 5A; Supplementary Table 7). GO analysis of significantly regulated genes from both groups highlighted pathways related to muscle system processes and cytokine-mediated immune responses (Supplementary Figure 5B). Within immune-related networks, two distinct regulatory patterns emerged (Supplementary Figure 4C). The first group included 21 immune-related genes, such as interleukins (*Il1rn*, *Il2rb*, *Il18rap*) and chemokines (*Ccl5*, *Ccl7*), which were significantly regulated in both STAT5B^Y665F^ and STAT5B^Y665H^ mutants in opposite directions. The second group consisted of 21 genes—including transcription factors (*Gata3*, *Stat4*, *Foxp3*), interleukins (*Il18ra*, *Il7r*, *Il12rb1*, *Il9r*), and chemokines (*Ccl22*, *Ccl8*, *Cxcr3*, *Cxcl14*, *Ccr5*, *Ccr7*)—that were significantly regulated in STAT5B^Y665F^ but not STAT5B^Y665H^ mice. These findings suggest that STAT5B^Y665F^ broadly activates lipid and immune transcriptional programs in adipose tissue, while STAT5B^Y665H^ exerts a more limited influence, highlighting their opposing and sex-dependent roles in systemic immune-metabolic regulation.

**Figure 6.**
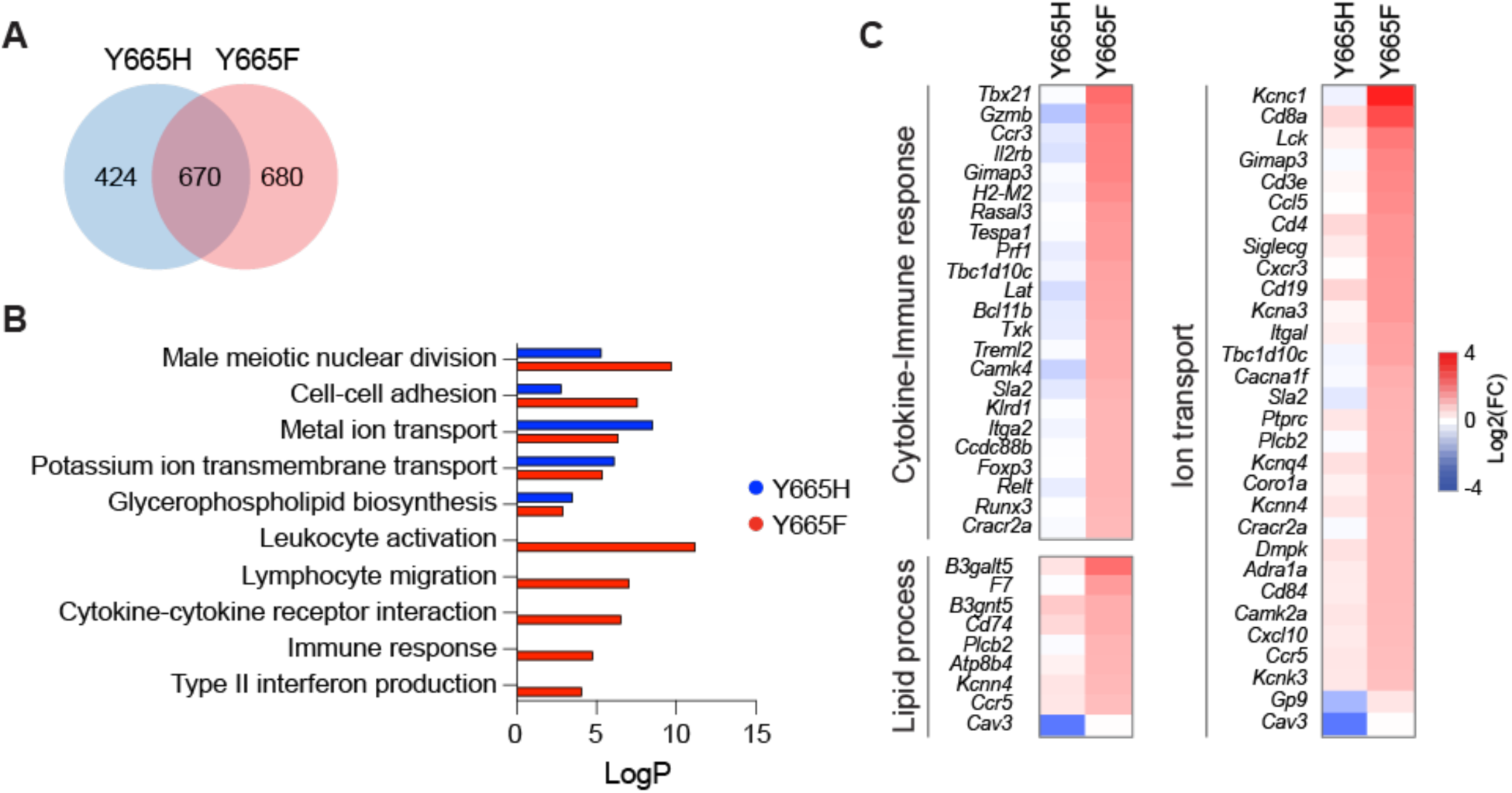
Opposing transcriptional activities of STAT5B^Y665F^ and STAT5B^Y665H^ in adipose tissue. **(A)** Venn diagram displaying significantly regulated genes in STAT5B^Y665^ mutant male mice compared to WT (RNA-seq, *n* = 6). **(B)** Bar graph of DEGs enriched in GO terms. **(C)** Heatmaps of log2(fold changes) for DEGs related to cytokine signaling in immune response, lipid metabolism and ion transport. Red, indicates genes induced compared to WT and blue indicates genes whose expression is reduced compared to WT.

## Discussion

Our study identifies Y665 as a critical regulatory residue within STAT5B, dictating opposing metabolic and transcriptional outcomes in vivo. Using CRISPR/Cas9 base editing to generate STAT5B^Y665F^ and STAT5B^Y665H^ alleles, we uncover a striking divergence in their functional impact on hepatic and systemic metabolism. STAT5B^Y665F^ acts as a potent gain-of-function variant, driving hepatic lipid accumulation, hypercholesterolemia, and enhanced insulin sensitivity, while STAT5B^Y665H^ exhibits a partial loss-of-function phenotype associated with reduced body mass and impaired insulin responsiveness. These findings underscore how subtle amino acid changes at a single residue can fundamentally reprogram the transcriptional output of STAT5B. Mechanistically, STAT5B^Y665F^ promoted widespread enhancer engagement and robust transcriptional activation of metabolic and immune pathways, consistent with its hypermorphic behavior. In contrast, STAT5B^Y665H^ was largely unable to bind chromatin or remodel enhancer landscapes, resulting in reduced transcriptional activity. Nevertheless, STAT5B^Y665H^ was able to partially feminize liver gene expression patterns. Notably, STAT5B^Y665F^ induced canonical cholesterol biosynthetic genes and immune regulators, linking STAT5B enhancer occupancy to systemic lipid metabolism and inflammatory signaling. This enhancer-centric mechanism highlights how STAT5B variants can function as master regulators of hepatocyte transcriptional identity.

The divergent metabolic phenotypes of STAT5B^Y665F^ and STAT5B^Y665H^ also reveal an unexpected duality: STAT5B^Y665F^ promotes lipid and cholesterol accumulation but concurrently enhances glucose clearance and insulin sensitivity, suggesting a tradeoff between lipid metabolism and glucose homeostasis. By contrast, STAT5B^Y665H^ appears to confer protection from lipid overload but at the cost of impaired insulin action. Such opposing outcomes may explain the spectrum of metabolic phenotypes associated with altered STAT5B signaling in humans; these include dyslipidemia, growth defects, and immune dysfunction. These findings align with previous studies demonstrating that STAT5 activity is indispensable for glucose and lipid homeostasis (1,43,44). STAT5 signaling in pancreatic β-cells is essential for insulin secretion and glucose tolerance, with activated STAT5 pathways improving glucose homeostasis and insulin sensitivity under dietary stress (45,46). Activating STAT5B mutations have been associated with increased insulin signaling and hyperinsulinemia, coupled with altered lipid profiles (47,48), which may account for the elevated cholesterol levels we observed in STAT5B^Y665F^ mice. Conversely, mice expressing a dominant-negative STAT5 variant exhibit impaired glucose tolerance, hyperinsulinemia, and leptin resistance (44,45), phenotypes consistent with the attenuated signaling seen in STAT5B^Y665H^ mice. Hepatocyte-specific deletion of STAT5 disrupts cholesterol homeostasis and enhances lipogenesis (6), reinforcing the role of STAT5 as a central node linking insulin and lipid signaling pathways.

Broader parallels can be drawn from models of other metabolic regulators. CRTC2 mutant mice, for example, display altered hepatic glucose production alongside global changes in cholesterol dynamics and insulin sensitivity (49,50). These interconnections are further supported by evidence that cholesterol accumulation impairs insulin signaling, while improved insulin sensitivity reciprocally modulates cholesterol metabolism (6,47). Similarly, independent lipid signaling pathways can regulate glucose metabolism, as seen in α/β2-syntrophin–null mice, where alterations in cholesterol and fatty acid handling reshape systemic glucose homeostasis (51–53). Together, these findings support a model in which hepatic STAT5B activation coordinates metabolic outcomes by synchronizing hepatic lipid metabolism, cholesterol balance, and insulin responsiveness.

Sex-specific effects observed in our study, further emphasize the complexity of STAT5B-mediated regulation. While STAT5B^Y665F^ elicited broad transcriptional reprogramming that substantially feminized the pattern of gene expression in male livers, its effects in female livers were more restricted, particularly within immune pathways. These differences reflect known sexual dimorphism in GH–STAT5 signaling (15,16,36). Additionally, the differential regulation of adipose tissue programs by STAT5B variants suggests a liver–adipose axis mediated by STAT5B-dependent transcriptional outputs. STAT5B^Y665F^ broadly activated immune and lipid networks in adipose tissue, whereas STAT5B^Y665H^ exerted limited control, highlighting their opposing and sex-dependent systemic roles.

Our observation that STAT5B^Y665F^ induces 77% of female-biased genes while repressing 51% of male-biased genes in male liver (Figure 3D) is reminiscent of our earlier finding that STAT5B variant N642H, a constitutively active form of STAT5 (STAT5B_CA_), induces widespread feminization of hepatic gene expression (41). Mechanistically, the sex-biased activity patters of hepatic STAT5B reflect the sex-differential temporal dynamics of pituitary GH secretion—pulsatile in males versus near-continuous in females—which drive corresponding patterns of STAT5B tyrosine phosphorylation, nuclear translocation, and chromatin binding (22,24,54). The similarity between transcriptional changes induced by STAT5B^Y665F^ and STAT5B_CA_ (41) in male liver, which share 361 upregulated genes enriched in immune and lipid metabolism pathways, supports the proposal that STAT5B^Y665F^ mimics the near-continuous activation of hepatic STAT5B characteristic of female liver (22,41). Further, our finding that STAT5B^Y665F^ shows extensive enhancer occupancy is consistent with the persistent chromatin accessibility expected from sustained STAT5 activity. Conversely, STAT5B^Y665H^ showed transcriptomic convergence with STAT5-deficient livers (14), consistent with its loss-of-function characteristics (32,33).

The mutations in STAT5B^Y665F^ and STAT5B^Y665H^ studied here are germline mutations. Consequently, an important question is the extent to which the observed hepatic phenotypes reflect changes in hepatocyte STAT5 activity *per se* versus indirect effects arising from changes in STAT5 activity in other tissues. STAT5 in adipose tissue regulates lipolysis and adipocyte differentiation (43,44), and our adipose tissue RNA-seq data revealed that STAT5B^Y665F^ broadly activates lipid metabolism networks while STAT5B^Y665H^ exerts more restricted effects, potentially affecting hepatic lipid accumulation through altered free fatty acid flux (42). Additionally, the elevated serum IGF-1 levels observed specifically in STAT5B^Y665F^ mice could alter the endogenous GH secretory pattern via negative feedback on pituitary GH secretion (55), potentially contributing to liver feminization independently of direct hepatocyte STAT5B^Y665F^ actions. Definitive determination of the relative contributions of hepatocyte-intrinsic versus systemic mechanisms would require hepatocyte-specific conditional knock-in models.

We made the striking observation that STAT5B^Y665F^ increased the expression of many female-biased genes in both male and female liver, to levels as high as 10- to 25-fold above those found in WT female liver. These increases were not associated with marked changes in STAT5 chromatin binding at or near the impacted genes, suggesting an indirect transcriptional mechanism. For example, increased cooperation of STAT5B^Y665F^ with other liver-enriched transcription factors or coactivators recruited to these loci, e.g., nuclear hormone receptors and histone acetyltransferases (13,56–59) may amplify the transcriptional output beyond what would be predicted from STAT5 binding alone. Alternatively, sustained activation of STAT5B^Y665F^ may more effectively displace transcriptional repressors such as BCL6, which normally restricts female-biased gene expression in male liver by competing with STAT5 for chromatin occupancy at their overlapping DNA motifs enriched nearby female-biased genes (24,60). STAT5-induced positive feedback loops or altered metabolite/hormone levels downstream of the induced genes could provide additional regulatory inputs. Distinguishing among these mechanisms will require analysis of STAT5 binding dynamics, identification of cooperating transcription factors through motif enrichment analyses, and characterization of chromatin architecture changes at these loci.

In summary, our findings establish STAT5B^Y665^ as a molecular switch that directs opposing enhancer and transcriptional programs, thereby reshaping systemic lipid and glucose metabolism. STAT5B^Y665F^ and STAT5B^Y665H^ represent contrasting models of STAT5B hyperactivation versus attenuation, offering mechanistic insight into how single-residue alterations rewire transcription factor function. Together with prior literature, our data place STAT5B at the nexus of insulin signaling, cholesterol homeostasis, and immune regulation, in addition to GH-regulation of liver sex differences. Given the prevalence of STAT5B mutations in human disease (61–64), these insights provide a conceptual framework for understanding how STAT5B activity influences metabolic health and may guide therapeutic strategies for dyslipidemia, insulin resistance, and GH/STAT5-related disorders.

## Materials and Methods

### Mice

Mice were housed and handled according to the Guide for the Care and Use of Laboratory Animals (8th edition). All animal experiments were approved by the Animal Care and Use Committee (ACUC) of National Institute of Diabetes and Digestive and Kidney Diseases (NIDDK, MD) and performed under the NIDDK animal protocol K089-LGP-20, K089-LGP-23 and K141-MMC-25. CRISPR/Cas9 and base editing (65,66) targeted mice were generated using C57BL/6N mice (Charles River) by the Transgenic Core of the National Heart, Lung, and Blood Institute (NHLBI). STAT5B^Y665H^ and STAT5B^Y665F^ mutant mice were germline point mutations generated using adenine base editing and CRISPR/Cas9 genome engineering methods, respectively, as described (32,33). Mice used in this study were heterozygotes for STAT5B^Y665F^ (activating mutation), as the homozygous mice could not be reliably aged for study due to development of an ulcerative dermatitis necessitating early euthanasia (32). STAT5B^Y665H^ (inactivating mutation) mice were homozygous. Two-month old male and female mice were used in the experiments, following ARRIVE guidelines (https://arriveguidelines.org/). WT littermate STAT5B^Y665H^ and STAT5B^Y665F^ mutant mice were used as controls. Tissues were collected from mice and used immediately or stored at -80°C.

### Metabolic phenotyping

Body composition (fat mass and fat-free mass) was measured in non-anesthetized mice using a time domain EchoMR-100H analyzer (Echo Medical Systems, Houston, TX). Glucose tolerance test (GTT, 2 g/kg, i.p.) was performed following an overnight fast. Insulin tolerance test (ITT, HumulinR, 0.75 unit/kg, i.p., Lilly, Indianapolis, IN) was conducted in non-fasted mice. Blood glucose was measured with a Glucometer Contour (Bayer, Mishawaka, IN) at the indicated time points in blood samples collected from the tail vein. For serum analyses, blood was collected at 10 AM from the tail vein of non-fasted mice. Free fatty acids (Fujifilm Waco Diagnostics, Mountain View, CA, reagents 999–34691, 995–34791, 991–34891, 993–35191), triglycerides (Pointe Scientific Inc., Canton, MI, T7532-120), cholesterol (Thermo Scientific, Middletown, VA, TR13421) were measured using the indicated colorimetric assays. Insulin and IFG1 was measured with an ELISA kit (Crystal Chem, Downers Grove, IL; ALPCO, Salem, NH). HOMA-IR (Homeostasis Model Assessment of Insulin Resistance) index was calculated as described (67).

### Total RNA sequencing (RNA-seq) and data analysis

Total RNA was extracted from frozen liver tissue of WT male and female (n=6 per group) and mutant (n=5-9 per group) mice using a homogenizer and the PureLink RNA Mini kit according to the manufacturer’s instructions (Thermo Fisher Scientific). The concentration and quality of RNA were assessed by an Agilent Bioanalyzer 2100 (Agilent Technologies, CA). Ribosomal RNA was removed from 1 μg of total RNA (RIN value > 8.0) and cDNA was synthesized using SuperScript III (Invitrogen). Libraries for sequencing were prepared from individual mice according to the manufacturer’s instructions with TruSeq Stranded Total RNA Library Prep Kit with Ribo-Zero Gold (Illumina, RS-122-2301) and paired-end sequencing was done with a NovaSeq 6000 instrument (Illumina).

Total RNA-seq read quality control (Supplementary Table 2) was carried out using Trimmomatic (68) (version 0.36) and STAR RNA-seq (69) (version 2.7.11b) using paired-end mode to align the reads to mouse genome release mm10). HTSeq (0.9.1) (70) was used to retrieve raw counts and subsequently, R (version 4.2.3) (https://www.R-project.org/), Bioconductor(71) and DESeq2 (72) were used. Additionally, the RUVSeq (73) package was applied to remove confounding factors. Data quality was assessed by PCA plot analysis. The data were pre-filtered, keeping only those genes that have at least 10 sequence reads in total. Genes were identified as significantly differentially expressed at log2 fold-change (FC) >1 or < -1 and adjusted p-value (pAdj) <0.05, to correct for multiple testing using the Benjamini-Hochberg method. Downstream analyses included gene enrichment analysis (GSEA, https://www.gsea-msigdb.org/gsea/index.jsp) and Metascape analysis (https://metascape.org/gp/index.html#/main/step1). Visualization was done using dplyr (https://CRAN.R-project.org/package=dplyr) and ggplot2 (74). Heatmaps were prepared with a custom heatmap helper package (https://github.com/Ye1203/Heatmap_shinyapp).

Liver RNA-seq data was also analyzed as described elsewhere (41) to directly compare the effects of each mutant form on genes that show significant differences in expression between male and female (WT) mouse liver. Results of these additional analyses are shown in Supplemental Table 3. These analyses include comparisons to gene responses induced in mice expressing the constitutively active STAT5B-N642H (STAT5B_CA_), a constitutively active STAT5B mutant whose expression substantially feminizes liver gene expression in male mouse liver (41). Briefly, sequence reads were aligned to mouse genome build mm9 (NCBI 37) using STAR aligner, and FeatureCounts was used to count sequence reads mapping to the union of the exonic regions of all isoforms of a given gene (collapsed exon counting). These analyses were based on an mm9 Gene Transfer Format file described and available elsewhere (75) and comprised of 75,798 mouse genes: n = 20,884 RefSeq protein coding genes, n = 48,363 mouse liver-expressed lncRNA genes, n = 2,061 RefSeq noncoding genes (NR accession numbers) that do not overlap the set of 48,360 lncRNAs, and n = 4,490 Ensembl noncoding lncRNAs that do not overlap either the RefSeq NR gene set or the 48,363 lncRNA gene set. Differential gene expression was determined with edgeR at log2 fold change >1 or <-1 and pAdj <0.05 to identify genes showing differences in expression between WT male and WT female liver. The impact of each STAT5B-Y665 mutation was determined based on pAdj <0.05. A set of 3,414 mouse genes was identified as stringently sex-independent in liver based on these filters: -1.15 < |linear fold-change| < 1.15, mean TPM (transcripts per million) > 5, and pAdj > 0.3 for differential expression between WT male and WT female liver.

Comparisons of the differentially expressed gene sets to the genes responsive to STAT5B_CA_ -treated male livers (41) and to hepatocyte-specific STAT5A/STAT5B-deficient livers (14) were made using our previously published datasets (Supplemental Table 1).

### Chromatin immunoprecipitation sequencing (ChIP-seq) and data analysis

Frozen-stored liver tissues were ground into powder in liquid nitrogen. Chromatin was fixed with 1% formaldehyde (Sigma-Aldrich) for 10 min at room temperature and then quenched with 0.125 M glycine for 10 min at room temperature. Nuclei were isolated using Farnham Lysis Buffer (5 mM PIPES pH 8.0, 85 mM KCl, 0.5% NP-40, supplemented with PMSF and proteinase inhibitor cocktails). The chromatin was fragmented to 250–500 bp using sonicator 3000 (30 cycles; 20 s pulse/20 s rest, Misonix Sonicators), checked on a DNA gel, and further lysed in RIPA buffer. One milligram of total protein was immunoprecipitated with Dynabeads Protein A (Novex) coated with ChIP-seq grade antibodies: STAT5B (R&D systems, AF1584; ThermoFisher scientific, 13-5300), H3K27ac (Abcam, ab4729) and RNA polymerase II (Abcam, ab5408). Libraries for next-generation sequencing were prepared with NEBNext Ultra II DNA library prep kit for illumina (New England Biolabs, M0287L) and sequenced with a NovaSeq 6000 instrument (Illumina).

Quality filtering and alignment of the raw reads was done using Trimmomatic (68) (version 0.36) and Bowtie (76) (version 1.3.1), with the parameter ‘-m 1’ to keep only uniquely mapped reads, using the reference genome mm10. Picard tools (version 2.9.2) (Broad Institute. Picard, http://broadinstitute.github.io/picard/ 2016) was used to remove duplicates and subsequently, Homer (77) (version 4.10.4) and deepTools (78) (version 3.5.4) software was applied to generate bedGraph files and normalize coverage, separately. Integrative Genomics Viewer (79) (version 2.3.98) was used for visualization. MACS2 (80) peak-finding algorithm was used to identify regions of ChIP-seq enrichment over the background, utilizing input files. For visualization, the total number of reads mapped in each sample was normalized to 10 million and background signals of <2 were eliminated. Each ChIP-seq experiment was conducted for two-three replicates and the correlation between replicates was computed by Pearson and Spearman correlation using deepTools.

### Histology and Immunohistochemistry

Liver tissues from WT and mutant mice were collected at two-months of age. Isolated tissues were fixed with 10% neutral formalin solution and dehydrated in 70% ethanol. Samples were processed for paraffin sections and stained with hematoxylin and eosin using standard methods (Histoserv). Images were captured using a Keyence BZ-9000 microscope.

### Statistical analyses

Samples used for RNA-seq were randomly selected, and blinding was not applied. For comparison of samples, data were presented as standard deviation in each group and were evaluated with a 1-way or 2-way ANOVA multiple comparisons using PRISM GraphPad (version 10.1.1). Statistical significance was obtained by comparing the measures from wild-type or control group, and each mutant group. A value of \**p* < 0.05, \*\**p* < 0.001, \*\*\**p* < 0.0001, \*\*\*\**p* < 0.00001 was considered statistically significant. ns, no significant.

### Data Availability

All data were obtained or uploaded to Gene Expression Omnibus (GEO). The RNA-seq and ChIP-seq generated in this study are available using accession code GSE310018; ChIP-seq under GSE309954, RNA-seq from liver tissue under GSE310017 and RNA-seq from adipose tissue under GSE309955. RNA-seq data from STAT5B_CA_ (41) and STAT5A/B-KO (14) livers were obtained from GSE196015 and GSE103885, respectively.

## Acknowledgements

This research was supported by the Intramural Research Programs (IRPs) of National Institute of Diabetes and Digestive and Kidney Diseases (NIDDK) within the National Institutes of Health (NIH). Work carried out at Boston University was supported in part by NIH grant DK121998 (to DJW). The contributions of the NIH author(s) were made as part of their official duties as NIH federal employees, are in compliance with agency policy requirements, and are considered Works of the United States Government. However, the findings and conclusions presented in this paper are those of the author(s) and do not necessarily reflect the views of the NIH or the U.S. Department of Health and Human Services.

We thank the NIH Intramural Sequencing Center (NISC) and the NHLBI sequencing core for next generation sequencing, and the NIDDK advanced light microscopy & image analysis core for microscopy equipment. This work utilized the computational resources of the NIH HPC Biowulf cluster (http://hpc.nih.gov) and Boston University Shared Computing Cluster.

## Author contributions

H.K.L. and L.H. initially designed the study and made initial data analyses. As the study progressed H.K.L., O.G., P.A.F., L.H. and D.J.W. contributed to further design of the study and data analysis.

H.K.L. established mutant mouse lines. H.K.L., N.L., T.D. and O.G. performed experiments. H.K.L., M.P., B.Y. and D.J.W. conducted data analysis. H.K.L., P.A.F, L.H. and D.J.W. wrote the manuscript, and all authors approved the final version. We acknowledge the use of ChatGPT (https://chat.openai.com/) to provide general information on topics discussed in this work and to clarify context.

## Ethical Approval

Mice were housed and handled according to the Guide for the Care and Use of Laboratory Animals (8th edition) and all animal experiments were approved by the Animal Care and Use Committee (ACUC) of National Institute of Diabetes and Digestive and Kidney Diseases (NIDDK, MD) and performed under the NIDDK animal protocol K089-LGP-20, K089-LGP-23 and K141-MMC-25.

## Figure legends

**Supplementary Figure 1.**
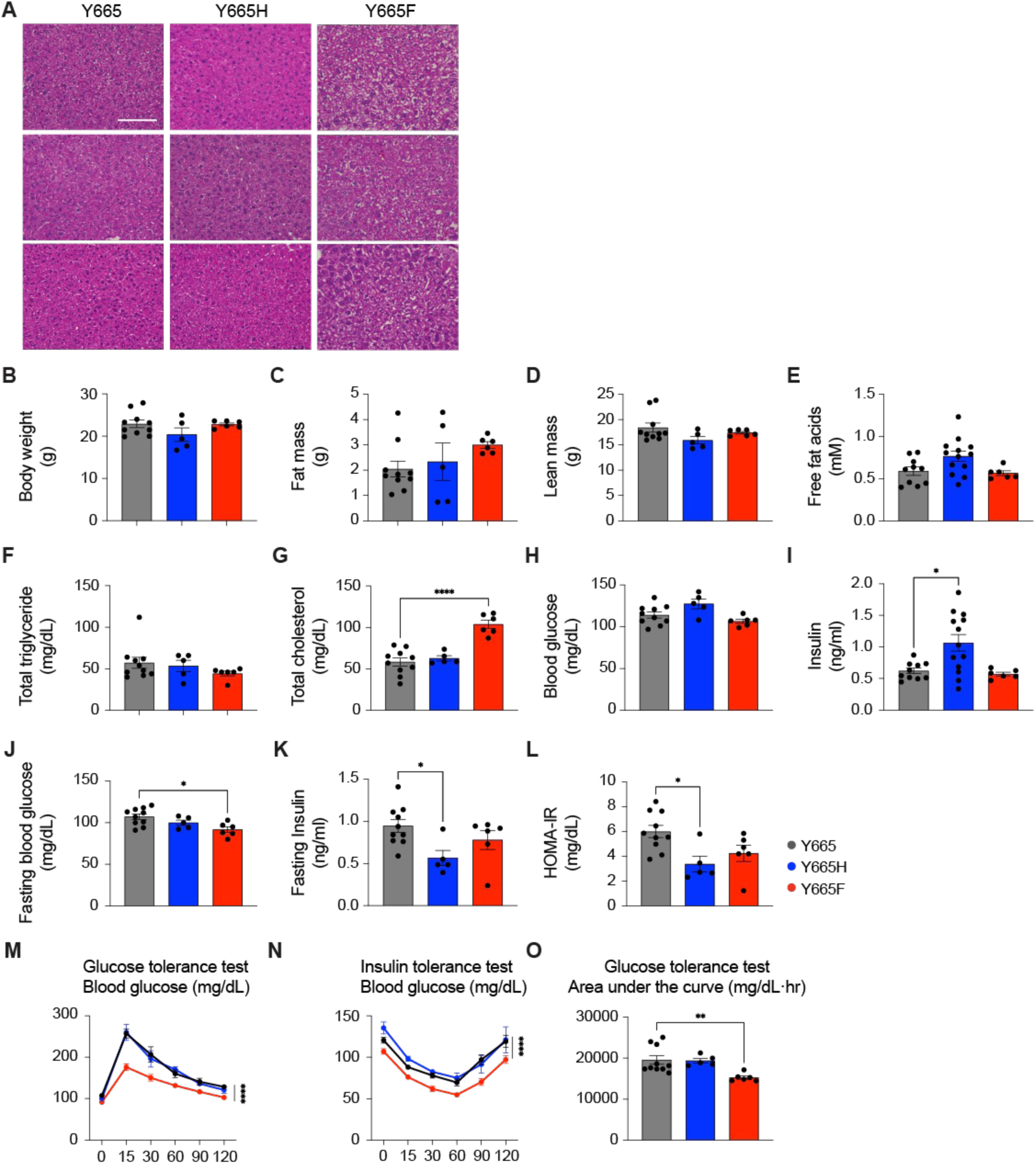
Sex-specific metabolic regulation by STAT5B^Y665^ mutations in female mice. **(A)** Representative hematoxylin and eosin (H&E)-stained liver sections from 2-month-old female mice (magnification x400; scale bars, 100 μm). **(B-I)** Physiological and biochemical parameters measured in 14-15-week-old WT and mutant female mice, including body weight (B), body composition (C-D), plasma free fat acids (E), total triglyceride (F), total blood cholesterol (G), blood glucose (H), and insulin (I). Data represent mean ± SEM (WT, *n* = 10; Y665H, *n* = 5; Y665F, *n* = 6). *P*-value were determined by one-way ANOVA with Tukey’s multiple comparisons test. \**p* < 0.05, \*\*\*\**p* < 0.0001. **(J-L)** Indices of insulin resistance and fasting state: fasting blood glucose (K), fasting insulin (L) and HOMA-IR (L). **(M-N)** Glucose and insulin tolerance tests (GTT and ITT) performed up to 120 min after glucose or insulin injection. *P*-value were determined by two-way ANOVA with Tukey’s multiple comparisons test. \*\*\*\**p* < 0.0001. **(O)** Glucose tolerance quantified by area under the curve (ACU). \*\**p* < 0.001.

**Supplementary Figure 2.**
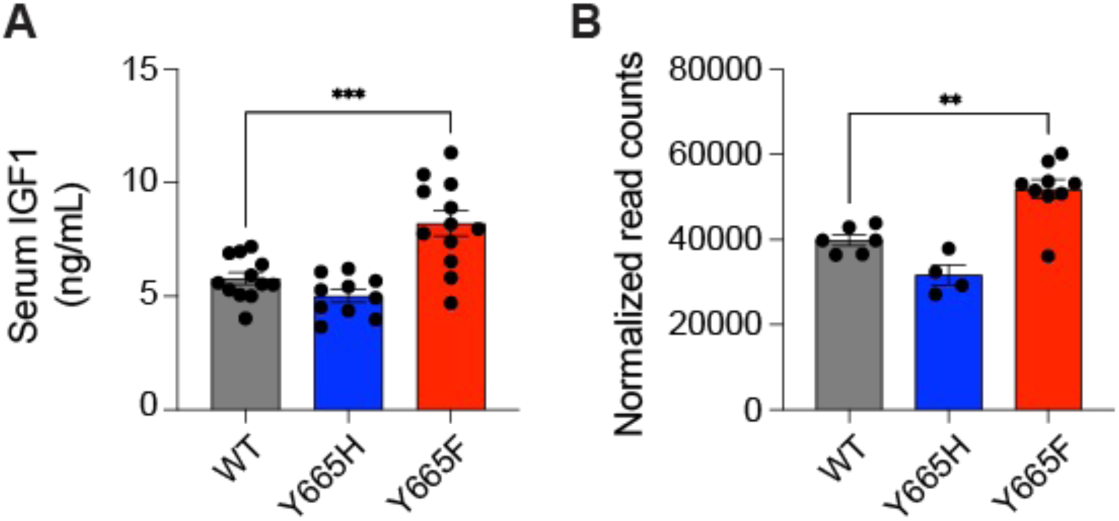
IGF-1 levels and its regulation in male STAT5B^Y665^ mutant mice. **(A)** Serum IGF-1 protein levels measured by ELISA. Data represent mean ± SEM (WT, *n* = 12; Y665H, *n* = 10; Y665F, *n* = 12). *P*-value were determined by one-way ANOVA with Tukey’s multiple comparisons test. \*\**p* < 0.001, \*\*\**p* < 0.0001. **(B)** Normalized RNA-seq read counts for *Igf1* (WT, *n* = 6; Y665H, *n* = 4; Y665F, *n* = 9).

**Supplementary Figure 3.**
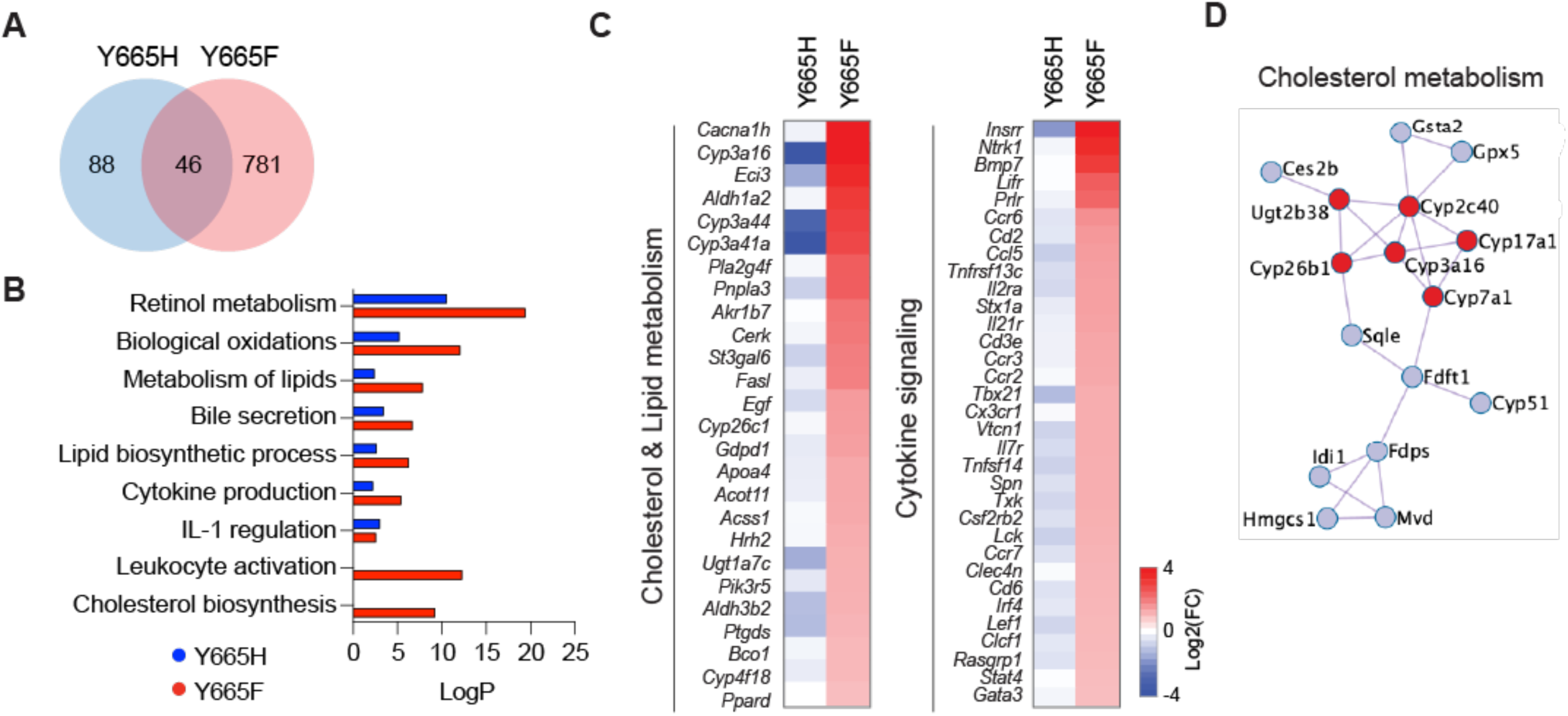
Transcriptional profiling of STAT5B^Y665^ mutants in female liver. **(A)** Venn diagram of DEGs identified by RNA-seq (*n* = 6). **(B)** GO term enrichment of female-specific DEGs. **(C)** Heatmaps of genes associated with cholesterol/lipid metabolism and cytokine signaling pathways. Red, indicates genes induced compared to WT and blue indicates genes whose expression is reduced compared to WT. **(D)** Cholesterol metabolism gene network reconstructed from RNA-seq data.

**Supplementary Figure 4.**
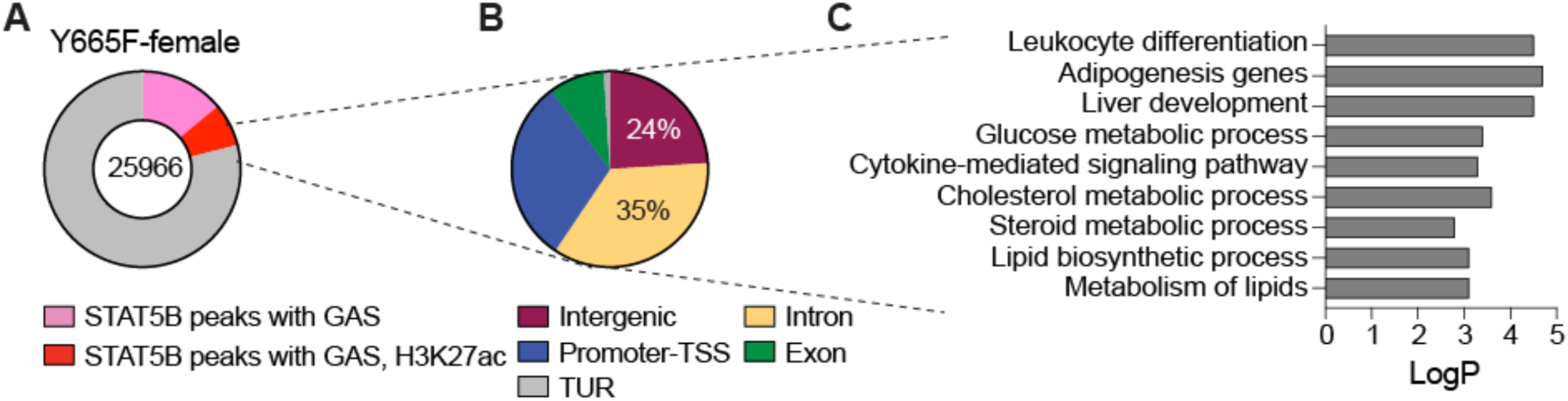
Enhancer landscape of STAT5B^Y665F^ mutants in female liver. **(A-B)** Distribution of STAT5B peaks containing GAS motifs and overlapping H3K27ac marks across genomic regions. **(C)** Functional annotation of enhancer-associated targets enriched in GO categories.

**Supplementary Figure 5.**
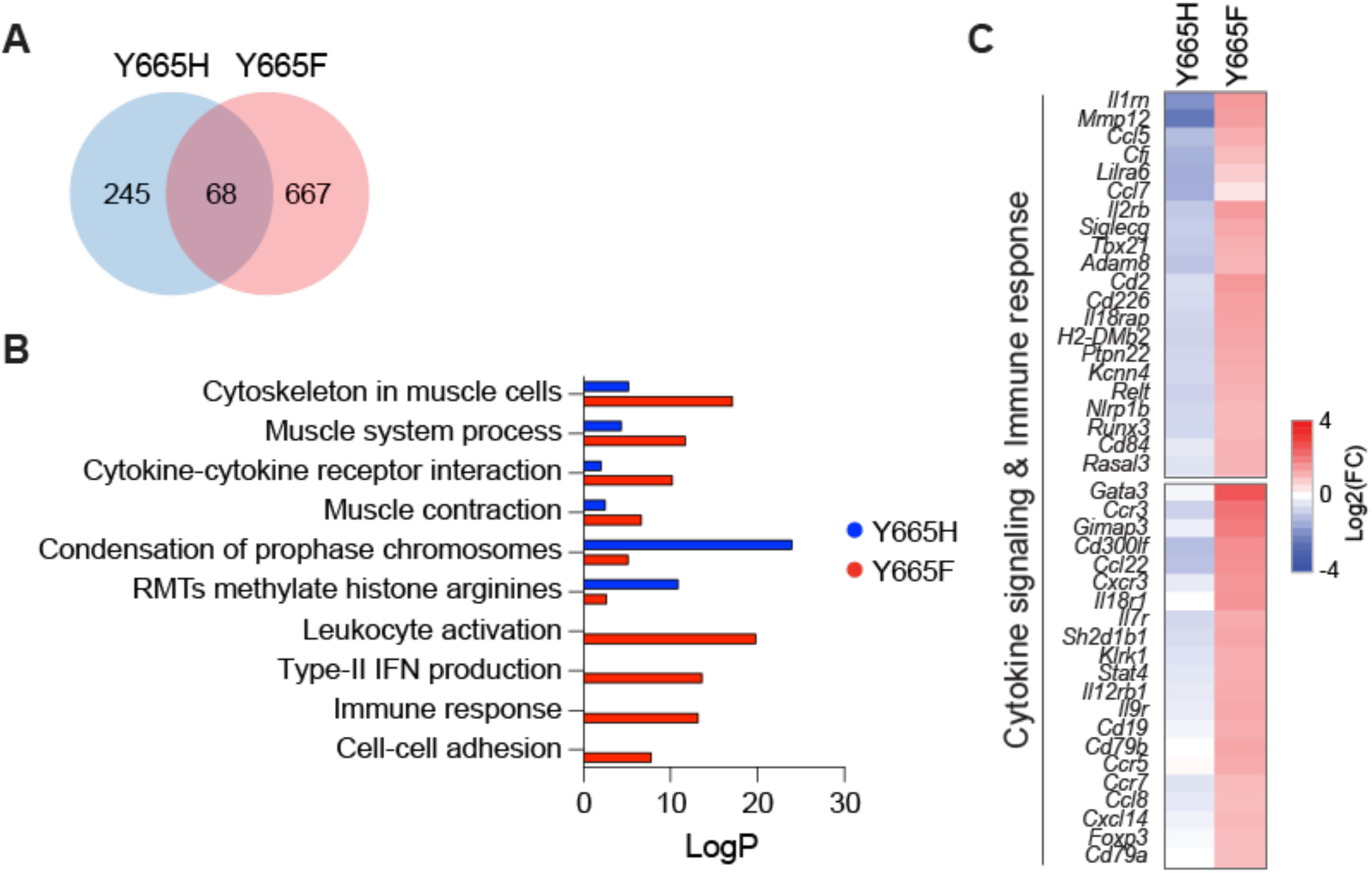
STAT5B^Y665^ mutations differentially reprogram transcription in female adipose tissue. **(A)** Venn diagram of DEGs from RNA-seq (*n* = 6). **(B)** Functional enrichment of DEGs in GO categories. **(C)** Heatmaps of immune- and cytokine-related DEGs highlighting mutation-specific effects. Red, indicates genes induced compared to WT and blue indicates genes whose expression is reduced compared to WT.

**Supplementary Table 1. Differentially expressed genes in male liver (STAT5B^Y665F^ and STAT5B^Y665H^ vs WT).** (A-B) Expression of significantly regulated genes: log2 (fold change), *p*-value, adjusted *p*-value and mean normalized read counts. (C-D) GSEA results for significantly regulated genes. (E-F) Comparison of transcriptomic responses in STAT5B^Y665^ mutant mice with those in STAT5B_CA_ or STAT5A/B-KO mice, respectively.

**Supplementary Table 2. Differentially expressed genes in female liver (STAT5B^Y665F^ and STAT5B^Y665H^ vs WT).** (A-B) Expression of significantly regulated genes: log2 (fold change), *p*-value, adjusted *p*-value and mean normalized read counts. (C-D) GSEA results for significantly regulated genes. (E) Comparison of transcriptomic responses in STAT5B^Y665^ mutant mice with those in STAT5A/B-KO mice.

Supplementary Table 3. Significantly regulated genes in male and female liver analyzed using GTF file representing 75,798 mouse genes. (A) Gene expression data. (B) Stringently sex-independent genes. (C-D) Expression data and cluster membership for genes shown in the heatmaps in Figure 4.

**Supplementary Table 4. Genome-wide STAT5B binding features in male liver.** A, C, E) STAT5B binding ChIP-seq binding peaks in WT and mutant male mice. (B, D, F) STAT5B peaks containing canonical GAS motifs (TTCnnnGAA) with corresponding H3K27ac coverage. (G) GO analysis of STAT5B-bound genes. (H) Expression levels of genes identified by ChIP-seq. Cholesterol and lipid metabolic genes are in red.

**Supplementary Table 5. Genome-wide STAT5B binding features in female liver.** (A) STAT5B ChIP-seq binding peaks in liver from STAT5B^Y665F^ female mice. (B) STAT5B binding peaks containing canonical GAS motifs (TTCnnnGAA) with H3K27ac coverage. (C) GO analysis of STAT5B-bound genes. (D) Overlap between STAT5B- and H3K27ac-associated genes.

**Supplementary Table 6. Differentially expressed genes in male adipose tissue (STAT5B^Y665F^ and STAT5B^Y665H^ vs WT).** (A-B) Expression of significantly regulated genes: log2 (fold change), *p*-value, adjusted *p*-value, and mean normalized read counts. (C-D) GSEA results for significantly regulated genes.

**Supplementary Table 7. Differentially expressed genes in female adipose tissue (STAT5B^Y665F^ and STAT5B^Y665H^ vs WT).** (A-B) Expression of significantly regulated genes: log2 (fold change), *p*-value, adjusted *p*-value, and mean normalized read counts. (C-D) GSEA results for significantly regulated genes.

